# Optimizing Tissue Clearing Methods for Improved Imaging of Whole-Mount Retinas

**DOI:** 10.1101/2025.04.14.648787

**Authors:** Aubin Mutschler, Volha V. Malechka, Petr Baranov, Jonathan R. Soucy

**Affiliations:** The Schepens Eye Research Institute of Massachusetts Eye and Ear, Boston, MA 02114; Department of Ophthalmology, Harvard Medical School, Boston, MA 02114; Program of Neuroscience, Middlebury College, Middlebury, VT 05753

## Abstract

Gene and cell therapies are promising approaches for restoring vision in hereditary and advanced optic neuropathies. However, these therapeutic approaches must be accurately evaluated through a combination of methods, including advanced imaging, to reach the clinic. We present a whole-mount tissue-clearing methodology to improve imaging of donor neuron integration in the retina following cell transplantation in mice. Mouse retinas were processed using five different clearing methods, comparing tissue transparency, pigmentation, and immunohistochemical clarity. Among the tested methods, ScaleS consistently outperformed its peers, demonstrating a 46% increase in tissue transparency and an 89% increase in immunohistochemical clarity compared to controls. We developed a modified version of ScaleS, termed ScaleH, by adding polyvinyl alcohol to reduce fluorescence decay and enhance sample stability. ScaleH maintained fluorescence stability over extended periods (32% less decay) and proved compatible with immunolabeling and endogenous fluorescent reporters, enabling improved visualization of transplanted human stem cell-derived retinal neurons in the mouse retina. Moreover, ScaleH also improved optic nerve imaging, demonstrating the potential for broader neurobiological applications. Our clearing workflow supports robust, high-resolution imaging for evaluating the integration of transplanted cells in regenerative studies.

## Introduction

As the global population ages, eye diseases such as glaucoma, age-related macular degeneration, and diabetic retinopathy are becoming increasingly prevalent, underscoring an urgent need for therapeutic advancements (Hashemi et al., 2017). Gene therapies and stem cell transplantation are promising approaches for restoring vision (Nanegrungsunk et al., 2022). The eye is particularly well-suited for these therapies due to its accessibility, immune privilege, and localized effects afforded by the blood-ocular barriers (Ghoraba et al., 2022). However, the precise targeting of therapies must be closely monitored. For example, retinal ganglion cell (RGC) transplantation requires integration into the ganglion cell layer and axon extension through the optic nerve to restore vision (Zhang et al., 2021)(Soucy et al., 2023). These challenges underscore the importance of robust models and advanced imaging techniques to refine and evaluate therapeutic approaches.

Understanding the molecular and cellular mechanisms underlying retinal function is essential for advancing our knowledge of ocular development and disease. Moreover, given its accessibility as part of the central nervous system, the retina offers a unique model for studying neurological processes beyond vision, with implications for broader biomedical applications (London et al., 2013).

Imaging is a vital tool for elucidating the interactions between novel treatments, molecular mechanisms, and the structural complexity of the retina. Optical coherence tomography (OCT) is the gold standard for non-invasive, three-dimensional imaging of the in vivo eye in humans and animals, yet it lacks the resolution needed to resolve the detailed cellular and molecular makeup of the retina (Huang et al., 1991)(Jagodzinska et al., 2017); thus, at this point, post-mortem histological assessment is needed to visualize subcellular structures. High-resolution approaches, such as single-photon confocal microscopy, enable the visualization of fine structures in thin tissue samples. In contrast, two-photon microscopy offers greater depth penetration but is limited by photobleaching and signal-to-noise challenges (Cheng et al., 2023). Alternatively, light-sheet fluorescence microscopy enables rapid, detailed three-dimensional imaging but often lacks the precision required for smaller structures, making confocal microscopy the preferred choice for retinal studies (Prahst et al., 2020).

Despite advances in imaging technology, optical limitations persist due to inherent tissue properties. Light scattering caused by structural heterogeneity and refractive index mismatches between lipid membranes and aqueous interfaces contributes to tissue opacity, limiting imaging depth and resolution, especially in thicker tissue samples (Richardson & Lichtman, 2015) (Chung et al., 2013). To overcome these obstacles, tissue-clearing techniques have been developed to reduce opacity (Richardson & Lichtman, 2015). Active clearing involves the use of external forces to remove lipids, requiring harsh reagents and complex protocols, while passive clearing utilizes chemical diffusion through aqueous or organic solvents. Hydrophilic-based solvents are preferred for fluorescent imaging due to their ability to preserve fluorescence and minimize tissue shrinkage (Renier et al., 2014). Recently, ultrafast clearing protocols have been developed to clarify tissue within minutes, reducing processing times and streamlining experimental workflows (Tomer et al., 2014)(Matsumoto et al., 2019). In facilitating high-resolution imaging, it is well-suited for cellular visualizations of the retina.

In this study, we compare five ultrafast tissue clearing protocols — ScaleS (Hama et al., 2015), FOCM (X. Zhu et al., 2019), OptiMuS (K. Kim et al., 2022), MACS (J. Zhu et al., 2020), and Visikol HISTO-1 (Villani et al., 2013) — to identify an optimal method for retinal applications. Our evaluation focused on three critical factors: tissue transparency, compatibility with immunostaining markers, and retention of fluorescence. ScaleS emerged as the optimal solution, and we developed a protocol tailored for retinal applications. To enhance its utility, we modified the ScaleS recipe to incorporate a self-hardening mechanism that maintains fluorescence over extended periods of time. Using this optimized protocol, we demonstrated compatibility with retinal tissue staining and visualization of stem cell-derived RGC transplantation in the retina. Lastly, we extended its application to the optic nerve, illustrating its versatility for other neural tissues beyond the retina.

## Methods and Materials

### Donor Cell Transplantation

All animal procedures were approved by the Institutional Animal Care and Use Committee (IACUC) at the Schepens Eye Research Institute. Donor RGC transplantations were performed as previously described (Soucy et al., 2023).

Donor retinal ganglion cells (RGCs) were differentiated from H9-BRN3B:tdTomatoThy1.2-hESCs using a 2D differentiation protocol (Sluch et al., 2017). Thy1.2-positive RGCs were isolated via magnetic microbead sorting, as previously described (Soucy et al., 2023). For transplantation, 20,000 RGCs suspended in 1 µL of N2B27 medium [1x GlutaMAX, 1x antibiotic antimycotic, 1% N2 supplement, and 2% B27 supplement in 50:50 1x DMEM/F12:Neurobasal medium] were injected intravitreally into the eyes of adult C57Bl/6J mice (4–8 months old, both sexes) under general anesthesia by intraperitoneal injection of ketamine/xylazine and local anesthesia (0.5% proparacaine eye drops). Injections were performed using a beveled glass microneedle (80 µm inner diameter) at a flow rate of 1 µL/min. Animals were maintained on a 12-hour light/dark cycle and euthanized 3 days post-transplantation via CO_2_ inhalation.

### Mouse Retina Sample Preparation

Retinal tissue and optic nerve samples were collected from C57Bl/6J wild-type mice aged 1-2 years of both sexes. Mice were euthanized by CO_2_ asphyxiation.

Following euthanasia, the eyes were enucleated, and the cornea was punctured (for whole mounts only) to improve the diffusion of the fixative. Samples were fixed in 4% paraformaldehyde (PFA; w/v in phosphate-buffered saline (PBS)) overnight on a shaker at 4°C.

PFA was prepared as frozen aliquots and thawed immediately before use. Eyes were washed with 0.01M PBS after fixation. For retinal whole mounts, retinas were then collected, and relaxing incisions were placed at equal distances around the retina for flat mounting. Isolated retinas were stored in 0.01M PBS overnight at 4°C for future use.

For retinal sections, whole eyes were sent to the Schepens Eye Research Institute Morphology Core for paraffin embedding (6 µm sections). To stain paraffin-embedded sections, they were first deparaffinized in xylene substitute I, washed with serial dilutions of ethanol, and then demasked for 30 min using sodium citrate buffer (10 mM Sodium citrate, 0.05% Tween 20, pH 6.0). To investigate an endogenous fluorophore signal, retinas from *Cx3cr1-GFP* mice were dissected, permeabilized, blocked, stained, cleared, and imaged within 12 hours.

### Clearing Solutions Preparation

OptiMuS, FOCM, ScaleSQ(5), and ScaleS4 were prepared as previously described (Kim et al., 2022) (Zhu et al., 2019) (Hama et al., 2015). In brief, to prepare the OptiMuS solution, 100 mM Tris (Sigma-Aldrich) and 0.34 mM EDTA (Invitrogen) were dissolved in ddH_2_O and titrated to a pH of 7.5 using 1N HCl, while being stirred continuously (200 rpm). After the buffer was prepared, 75% (w/v) Histodenz (Sigma-Aldrich) was added and dissolved at 60°C with continuous stirring (200 rpm). Subsequently, 10% (w/v) D-sorbitol (Sigma-Aldrich) and 4M urea (Sigma-Aldrich) were dissolved into the solution under the same conditions (60°C, 200 rpm). The completed OptiMuS solution was cooled to room temperature and stored at 4°C for later use.

To prepare the FOCM solution, 30% (w/v) urea and 20% (w/v) d-sorbitol were dissolved in DMSO with continuous stirring (200 rpm) overnight at room temperature. After complete dissolution, glycerol (Sigma-Aldrich) was added to the solution while stirring continued. The FOCM solution was stored at room temperature for later use.

To prepare the ScaleSQ(5) solution, 22.5% (w/v) D-(–)-sorbitol and 9.1M urea were dissolved in ddH_2_O with continuous stirring (200 rpm). Subsequently, 5% (w/v) Triton X-100 was added and stirred until fully dissolved. The solution was titrated to a pH of 8.2. The complete ScaleSQ(5) solution was stored at room temperature for future use.

To prepare ScaleS4, 40% (w/v) D-(–)-sorbitol, 10% (w/v) glycerol, 4M urea, and 20% (v/v) dimethyl sulfoxide (FisherSci) were dissolved in ddH_2_O and titrated to a pH of 7.8. Subsequently, 0.2% (w/v) Triton X-100 was added, and the solution was stored at room temperature for future use.

Lastly, to prepare our custom ScaleH combination clearing and mounting solution, two solutions (A and B) were combined at 55°C. Solution A was prepared by dissolving 15% (w/v) polyvinyl alcohol and 15% (w/v) glycerol in deionized water, followed by overnight incubation at room temperature with rocking. The solution was subsequently heated at 55°C until clear. Solution B was prepared by dissolving 3 M urea, 22.5% (w/v) D-(–)-sorbitol, 0.1 M Tris-HCl, 2.5% (w/v) 1,4- Diazabicyclo[2.2.2]octane (DABCO), and 15% (v/v) DMSO in deionized water at 55°C with continuous stirring. Once both solutions were warm and fully dissolved, Solution A was added to Solution B with stirring until homogenized. The pH was adjusted to 8.7 using HCl or NaOH. The final ScaleH solution was aliquoted into vials and stored at –20°C for future use.

### Immunohistochemistry

For immunofluorescent staining of whole-mount retinas and retinal sections, tissue were permeabilized using respective protocols. Samples were incubated in FOCM permeabilization (30% (w/v) sucrose in 0.01M PBS) and ScaleS permeabilization (20% (w/v) sucrose in 0.01M PBS) solutions for 4 hours at 4°C or the OptiMuS permeabilization (20% (w/v) DMSO and 2% (w/v) Triton X-100 in 0.01M PBS) solution for 4 hours at 37°C and subsequently washed three times with 0.1% triton X-100 and 0.1% tween-20 in 0.01M PBS (PBST) in 10-minute intervals. Permeabilization and wash steps for Militenyi Biotec^TM^ MACS® and Visikol® HISTO™ were completed following the Militenyi Biotec^TM^ mouse brain hemispheres and Visikol® protocols. Control samples were washed three times with PBST in 10-minute intervals.

Retinas were then blocked against non-specific antibody binding using the OptiMuS blocking buffer (10% (w/v) bovine serum albumin in 0.01M PBS), the Visikol® HISTO™ blocking solution, or our custom blocking buffer (10% goat serum, 1% bovine serum albumin, 0.1% sodium citrate, 0.25% tween-20, 0.25% triton-X, and 0.3M glycine in 0.01 M PBS) as previously described (Soucy et al., 2023). Samples were incubated for 2 hours at room temperature.

Following blocking, the tissue was incubated with primary antibodies in each clearing solution’s corresponding staining buffer. For OptiMuS, FOCM, ScaleS, and our control, the staining buffer consists of 1% bovine serum albumin, 0.25% Tween-20, and 0.25% Triton-X in 0.01M PBS. Staining buffers for MACS® and HISTO™ were prepared according to the manufacturer. A complete list of primary antibodies and their concentrations used in this study is available in **Table S1**. All retinal whole-mounts were incubated with primary antibodies for 60 hours at 4°C. Retinal sections were incubated overnight at 4°C. Samples were washed three times in 1-hour intervals using PBST or the manufacturer’s specified wash buffer. After removing any unbound primary antibody, goat anti-chicken, anti-guinea pig, anti-rabbit, and anti-mouse secondary antibodies (Jackson Immuno) were prepared at 1:500 in each staining buffer and incubated for 24 hours at 4°C. Samples were then washed with wash buffer in triplicate to remove any unbound secondary antibody and then incubated with 1 µg/mL DAPI (4’,6-diamidino-2- phenylindole) in PBS for 1 hour for whole mounts and 20 min for sections to stain the cell nuclei. For the MACS and Visikol tissue clearing, samples were dehydrated according to each manufacturer’s protocols.

### Imaging

Prior to clearing, retinal tissues were visualized using a stereomicroscope to verify flat-mount placement on single concave microscope slides. To evaluate pigmentation, images of each retina before and after clearing were taken with a white background, illuminated uniformly by an LED ring light (Unitron). Images were captured using a 4K Digital Wi-Fi Microscope (Jiusion-4KScope) positioned within the ring light. One retina from each group was placed above the Harvard Medical School logo and imaged to qualitatively visualize sample transparency.

To quantify transparency, retinas before and after clearing were imaged on top of a Fine Science Tools ruler with clearly delineated demarcation lines using an Olympus BX63 microscope with Olympus cellSens Dimension software and the Olympus XM10 camera. Imaging conditions were standardized, including consistent ambient lighting, LED brightness set at 0, exposure time of 174.7 ms, and gain of 8 dB. Images were captured at 2x magnification (1376 × 1038 pixels).

Samples were prepared for clearing by drying them using an absorption spear and applying 60µL clearing solution onto the microscope slides. The standard clearing protocol utilized a 10- minute incubation, with ScaleS clearing requiring an additional 10-minute ScaleS4 solution incubation post-ScaleSQ treatment. Post-clearing transparency imaging replicated pre-clearing conditions, including ruler superimposition and standardized background illumination. Samples previously imaged with institutional logos were imaged identically post-clearing.

Subsequently, samples were positioned coverslip-down for imaging with an Olympus FLUOVIEW FV3000 confocal microscope. Flat retinal regions were selected for imaging using one-way 1024 × 1024 Galvano scanning with frame intervals and step sizes set to 2 µm. Preliminary visualization at 2x magnification was used to determine precise areas of interest, which were subsequently imaged at 20x magnification. Following imaging, samples were stored at room temperature under minimal light exposure to preserve integrity.

### Image Analysis

Images were loaded into ImageJ and resliced from the left-hand side, with output rotations applied for consistent orientation. Intensity profile plots for each slice were generated, and data was exported as text files. Results were cleared for subsequent slices.

A custom MATLAB script was then used to analyze the previously generated cell nuclei intensity profile plots. Data files containing slice information were batch-imported and processed using the MATLAB trapz function to calculate the integral of intensity curves for each image slice.

Integrals were normalized by dividing by the number of data points along the x-axis. The first local maximum for each slice was measured, and the largest first local maximum across all slices in a retina was selected to normalize the integrals. Normalized integrals were averaged for each retina.

Microglial analyses focused on flat retinal sections, excluding unsuitable images. Images underwent background subtraction in ImageJ, followed by left-hand side reslicing and 90-degree rotation. A maximum-intensity z-projection was applied, and intensity profiles were generated for analysis. Coordinates corresponding to peak microglial intensities were identified and normalized to a value of 1. Slopes between two peak intensity points were calculated for each clearing solution, with average x and y coordinates recorded for comparative plotting.

## Statistical Analysis

Statistical significance was calculated using GraphPad Prism 10, employing an ordinary one-way ANOVA. Data are presented as mean ± standard deviation (*P < 0.05, **P < 0.01, ***P < 0.001, and ****P < 0.0001). Each data point within the bar plots represents one flat-mount retina.

## Figure Generation

Figures were generated using Adobe Illustrator 2024, Adobe Photoshop 2024, BioRender, and GraphPad Prism 10.

## Data/Code Availability

A custom MATLAB script used to measure normalized DAPI intensity (Fig. 2C) is available upon request.

**Figure 1.**
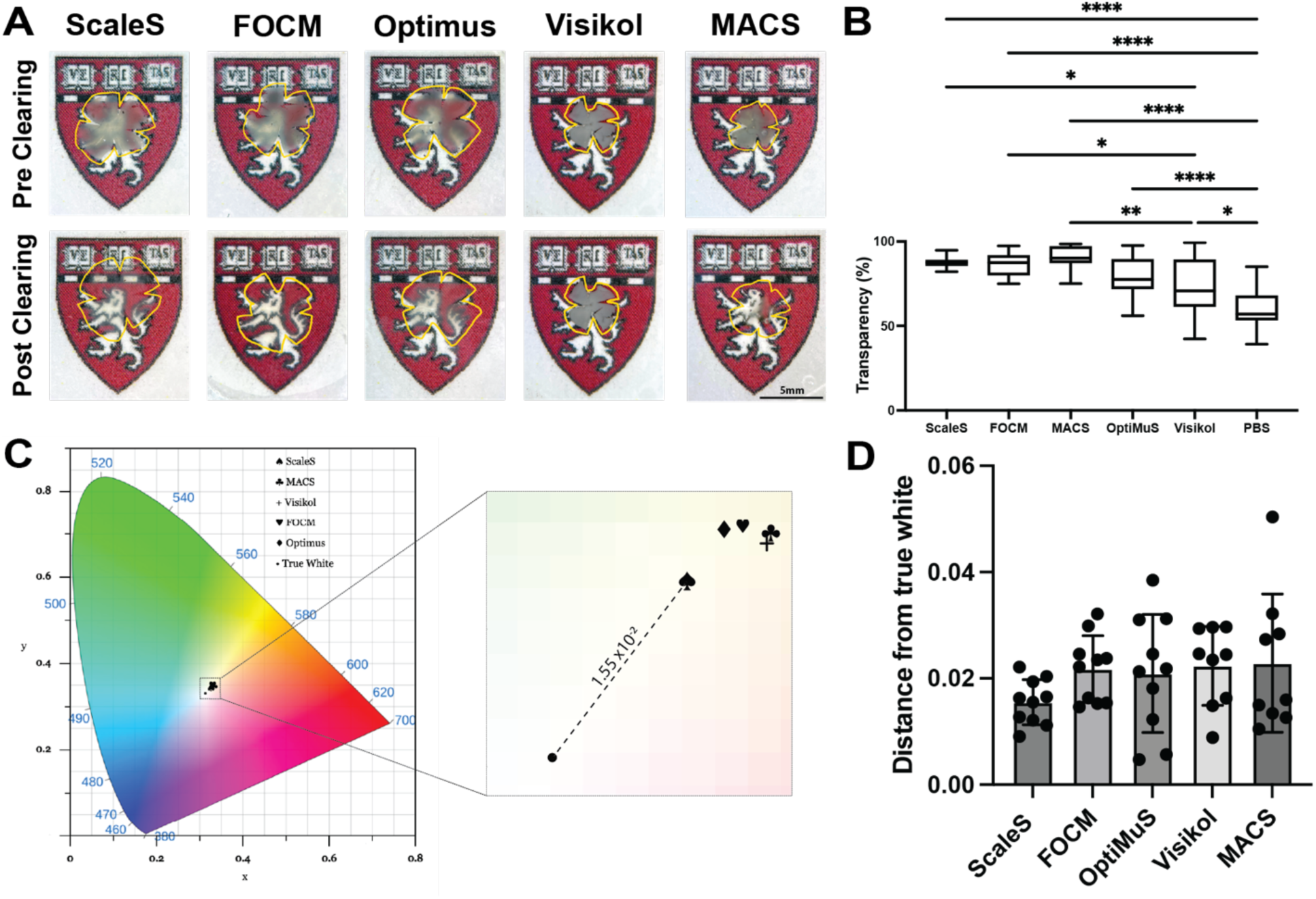
Evaluation of optical transparency and color across retinal clearing methods. **(A)** Representative en face images of whole-mount retinas before and after clearing with ScaleS, FOCM, OptiMuS, Visikol HISTO-1, and MACS, overlaid on the Harvard Medical School logo to visualize transparency improvements. Yellow outlines indicate the retina boundary. **(B)** Quantification of tissue transparency, calculated as the difference in grayscale value ratios between obstructed and unobstructed lines pre- and post-clearing. One-way ANOVA (F(5, 93) = 28.1, p < 0.0001) with Tukey’s post hoc test; *p < 0.05, **p < 0.001, ***p < 0.0001, ****P < 0.0001. N = 10-12 retinas per condition; N = 43 for PBS control. **(C)** CIE 1931 chromaticity diagram showing perceived color shift of cleared tissues relative to true white, based on x and y coordinates from color space transformations. Inset highlights the distance from the true white reference. **(D)** Quantitative assessment of color fidelity, defined as the Euclidean distance between each clearing method’s average color coordinates and the true white point. N = 9–10 retinas per group.

**Figure 2.**
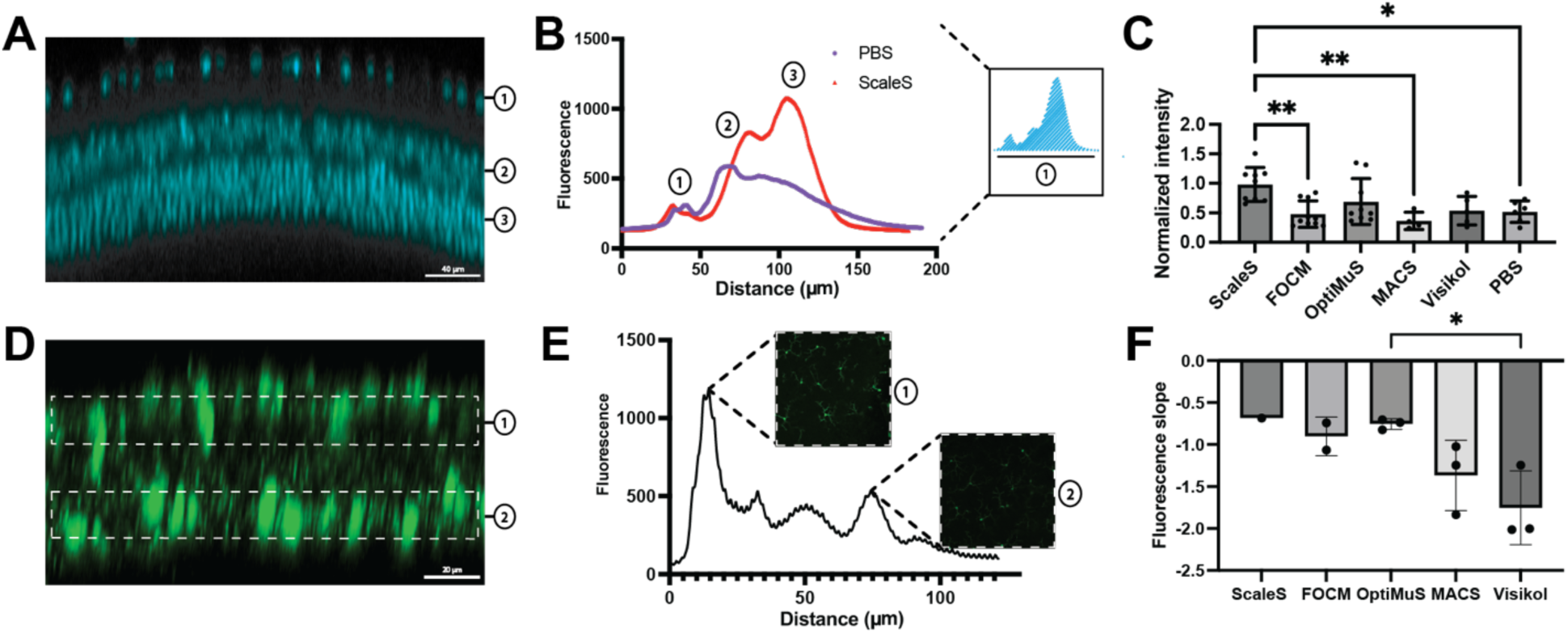
Quantitative comparison of clearing agents based on nuclear and microglial fluorescence retention. **(A)** Representative immunofluorescence cross-section of whole-mount retina (max intensity projection) stained for DAPI with labeling of (1) retinal ganglion cells, (2) bipolar cells, and (3) photoreceptors. **(B)** Representative fluorescence intensity profiles across depth for DAPI staining in control (PBS) and ScaleS-cleared samples show enhanced intensity and depth for ScaleS with quantification analysis inset. **(C)** Quantification of normalized nuclear fluorescence, defined as the average of integrals over the largest first local maximum, across clearing agents. N = 4-10 retinas per group. **(D)** Representative cross-sectional images of Iba1+ microglial layers in the retina. **(E)** Depth-resolved fluorescence intensity of Iba1 signal shows loss across the z-axis. Insets show representative en face microglial morphology within the two microglial layers. **(F)** Quantification of fluorescence slope between microglial layers for each clearing agent. N = 1-3 retinas per group. *p < 0.05.

## Results

### Clearing Improves Transparency of the Retina

To identify the most effective tissue-clearing method for retinal tissue, we evaluated five established clearing solutions based on immunohistochemical compatibility, ease of use, and demonstrated clearing efficacy: ScaleS (Hama et al., 2015), FOCM (X. Zhu et al., 2019), OptiMuS (K. Kim et al., 2022), MACS (J. Zhu et al., 2020), and Visikol HISTO-1 (Villani et al., 2013). Each solution was applied to mouse retinas, and transparency was assessed qualitatively and quantitatively.

In preliminary assessments, all solutions except Visikol HISTO-1 qualitatively increased retinal transparency (Fig. 1A). Post-clearing, tissues treated with MACS and Visikol became notably denser and more brittle. Transparency improvements were quantitatively assessed by measuring percentage changes in pixel gray values along defined reference lines (Fig. 1B; SI Appendix, Fig. S1). We observed significant differences in transparency across clearing methods (one-way ANOVA, F(5, 93) = 28.1, p < 0.0001). Tukey’s post hoc test showed that ScaleS was more transparent than Visikol (p = 0.0247) and PBS control (p < 0.0001), FOCM was more transparent than Visikol (p = 0.0272) and PBS control (p < 0.0001), OptiMuS and MACS were both superior to PBS control (p < 0.0001), MACS was better than Visikol (p = 0.0022), and Visikol outperformed PBS control (p = 0.0130). FOCM, ScaleS, and MACS increased transparency by 44%, 46%, and 50% compared to the control, with ScaleS showing the most consistent improvements in transparency.

In addition to transparency, we also evaluated the removal of pigmentation. Tissue pigmentation can cause severe light absorption and reduce image quality (Zhu et al., 2021). To quantify pigmentation removal, we developed a novel analysis method using a color space map (Zhu et al., 2019)(Hasabeldaim et al., 2023). Using chromaticity diagrams to map color perception, we converted RGB data to the XYZ color space to measure hue differences resulting from each clearing solution (Fig. 1C; SI Appendix, Fig. S2). The distance between each solution’s chromaticity value and a true white reference point was calculated to approximate pigmentation clearance (Fig. 1D).

Although differences in chromaticity distances were not statistically significant, ScaleS achieved the shortest distance to true white, achieving a color value 26% closer to true white than the nearest competitor. Furthermore, it displayed minimal variation compared to other solutions, underscoring its consistent performance. Together, these findings identified ScaleS, OptiMuS, MACS, and FOCM as promising solutions for enhancing retinal transparency.

### Tissue Clearing Enhances Full-thickness Immunostaining of the Retina

After evaluating transparency, we quantified the clearing effectiveness of each solution by comparing fluorescence intensity across retinal layers. DAPI staining, which labels the nuclei of cells in three distinct retinal layers, served as a benchmark for interlayer comparison (Fig. 2A). The RGC layer, positioned superficially, was used as a reference for normalization. Bipolar and photoreceptor cell layers, located progressively deeper within the retina, were analyzed to assess tissue transparency.

To quantify DAPI fluorescence penetration, we employed a semi-automated analysis of intensity profiles (Fig. 2B). Integrated fluorescence intensity across retinal layers was calculated and normalized relative to the brightest signal in the RGC layer of each retina. Statistical analyses of normalized intensities revealed that ScaleS (0.98) produced significantly higher staining intensities of the nuclei in the retina. Signal intensity was used as a proxy for clearing efficacy, especially within the deepest layer, as the nuclear signal from the photoreceptor layer would be attenuated as one moved deeper into the tissue. In contrast to the other clearing solutions tested, FOCM (0.48) and MACS (0.36) produced lower fluorescence intensities, indicating less effective tissue clearing. The reduced signal suggests limited light penetration and decreased visibility of deeper retinal layers, indicating lesser transparency than ScaleS (Fig. 2C).

In addition to nuclear staining, we assessed fluorescence penetration through microglial labeling. Retinal microglia are arranged in two distinct layers, allowing us to evaluate signal attenuation between superficial and deeper layers using slope analysis (Fig. 2D) (Fig.S3). Peak intensity values from fluorescence profiles were selected to calculate slopes, with a slope closest to zero representing no fluorescence attenuation throughout the retina (Fig. 2E).

Through this analysis, we showed the average slopes of ScaleS (m=-0.68) and OptiMuS (m=- 0.75) were the closest to zero, indicating limited signal degradation through retinal tissue and the greatest tissue transparency (Fig. 2F). For samples that exhibited poor tissue transparency, the discrepancy between microglial layers was larger, corresponding to slopes of m=-1.4 and m=-1.8 for MACS and Visikol, respectively.

### Development of a Hard-setting Clearing Solution (ScaleH) to Enhance Fluorescent Stability in Application Within and Beyond the Retina

Building upon the demonstrated efficacy of ScaleS, we developed a modified clearing formulation, termed ScaleH, by incorporating polyvinyl alcohol (PVA) to produce a hard-setting medium capable of preserving fluorescence signals in cleared tissues. While commercial hard- setting mounting media, such as ProLong Glass antifade mountant, can limit fluorescence signal decay, they are not optimized for mounting cleared tissue. ScaleH was designed to minimize signal decay over time, allowing for prolonged visualization in cleared tissue samples.

To evaluate ScaleH for compatibility with immunostaining (Fig. 3A), we stained whole retinal flat mounts for key retinal neuron populations. Specifically, photoreceptors, bipolar cells, and retinal ganglion cells were labeled using recoverin, PKC-α, and Brn3a, respectively (Hoon et al., 2014). Samples were counterstained with DAPI to visualize the cell nuclei. Computationally reconstructed sections of cleared retinal flat mounts showed clear visualization of all cell layers and nuclei in all experimental conditions (Fig. 3B). En face imaging confirmed specific staining of photoreceptor outer segments, bipolar cells, and retinal ganglion cells (Fig. 3B).

**Figure 3.**
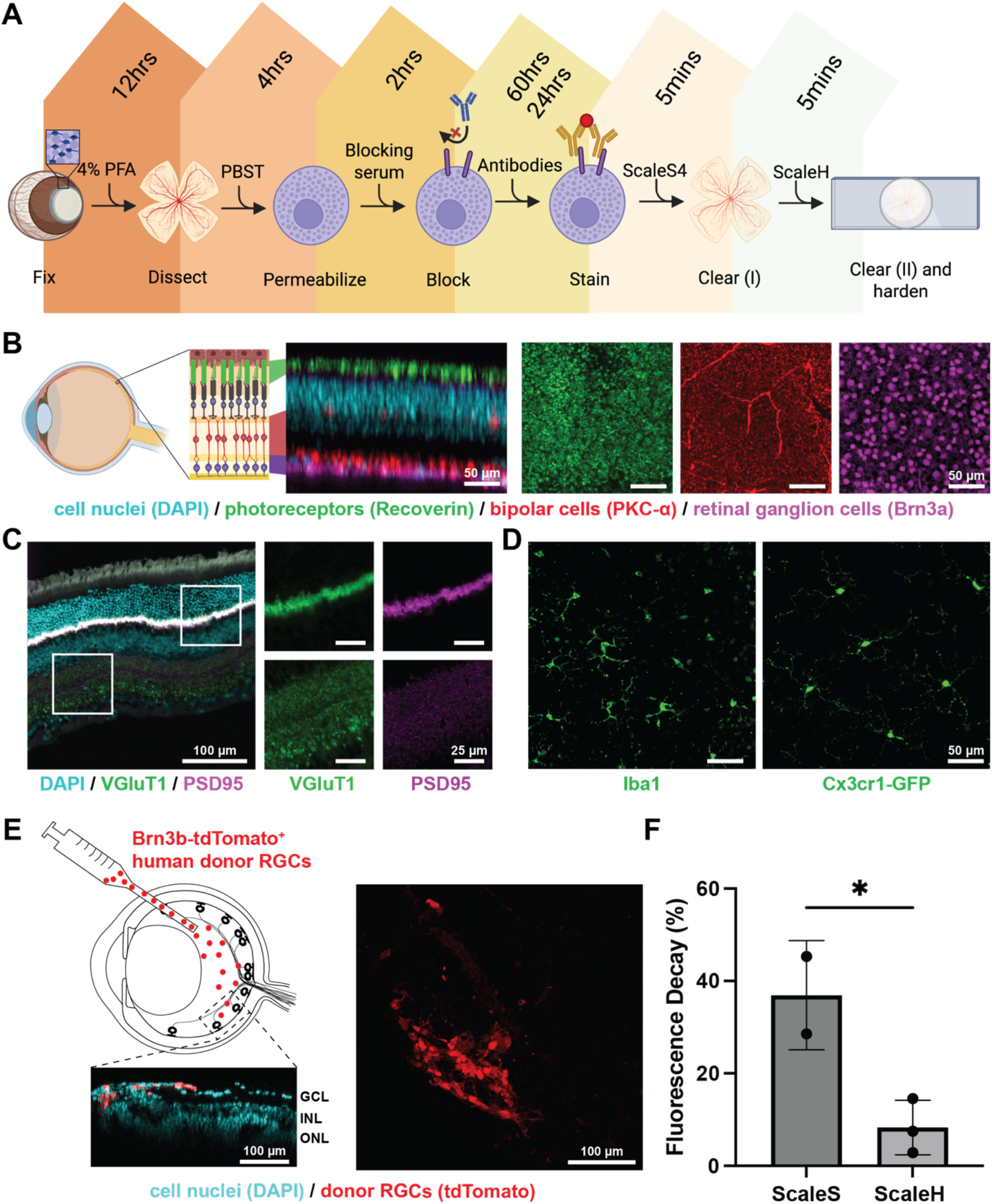
Self-hardening ScaleH is compatible with retinal imaging and improves the long-term preservation of fluorescence. **(A)** Workflow schematic of retinal tissue clearing using the self-hardening ScaleH protocol: fixation, permeabilization, blocking, antibody staining, and sequential clearing with ScaleS4 followed by ScaleH. **(B)** Representative immunofluorescent orthographic projection and en face images of a retinal whole mount successfully labeled with DAPI for cell nuclei, Recoverin for photoreceptor outer segments, PKC-α for bipolar cells, and Brn3a for retinal ganglion cells after tissue clearing. **(C)** Representative immunofluorescent images of retinal sections stained for DAPI, VGluT1, and PSD-95 after tissue clearing with the ScaleH protocol. **(D)** Representative immunofluorescence en face images of retinal microglia visualized by Iba1 staining and Cx3cr1 GFP-tagged microglia after clearing show that ScaleH is compatible with fluorescent reporters. **(E)** Representative immunofluorescence images show transplanted human stem cell-derived retinal ganglion cells (RGCs, red) in the mouse retina, imaged after clearing with ScaleH. Inset shows an orthographic projection with DAPI (blue) marking donor and host cell nuclei. GCL, ganglion cell layer; INL, inner nuclear layer; ONL, outer nuclear layer. Maximum intensity projection shows donor RGCs labeled with tdTomato. **(F)** Quantification of fluorescence preservation over two weeks shows significantly reduced signal decay in ScaleH-treated samples compared to ScaleS. N = 2-3 retinas per group. *p < 0.05.

Next, we evaluated the preservation of synaptic markers, which is crucial for examining neural connectivity and identifying synaptic pathology in retinal disorders (Graham & Duan, 2020).

Immunostaining of retinal sections with synaptic markers VGLUT1 and PSD-95 resulted in correctly localized staining patterns (Fig. 3C).

We also examined compatibility with endogenous fluorescent reporter expression using microglia-specific heterozygous *Cx3cr1^GFP/+^* mice. Microglia are essential for immune surveillance, synaptic pruning, and neuroinflammation in the retina (Ramirez et al., 2017). Immunostaining with the microglial marker Iba1 confirmed robust labeling across all clearing protocols. Importantly, clearing procedures did not disrupt endogenous GFP fluorescence in microglia (Fig. 3D).

To assess ScaleH performance in experimental models, we stained retinas containing human stem cell-derived RGCs post-transplantation. Donor RGCs were clearly visualized within appropriate retinal layers (Fig. 3E), with strong DAPI and tdTomato signals. Quantification of fluorescence signal over two weeks by the same approach described above revealed significantly reduced signal decay in ScaleH-treated samples (8%) compared to ScaleS (40%) (Fig. 3F).

### ScaleH Enables Clearing of the Optic Nerve

To evaluate the broader applicability of ScaleH, we applied the clearing protocol to mouse optic nerves, which are densely myelinated and structurally compact. These properties typically limit optical imaging depth in uncleared samples. Following treatment with ScaleH, optic nerves displayed visibly increased transparency (Fig. S4). En face and orthographic images of DAPI-labeled nuclei showed clear, continuous labeling throughout the entire cross-sectional area of the nerve, whereas samples mounted in ProLong Glass exhibited DAPI signal primarily at the surface, with limited penetration into deeper regions (Fig. 4A).

**Figure 4.**
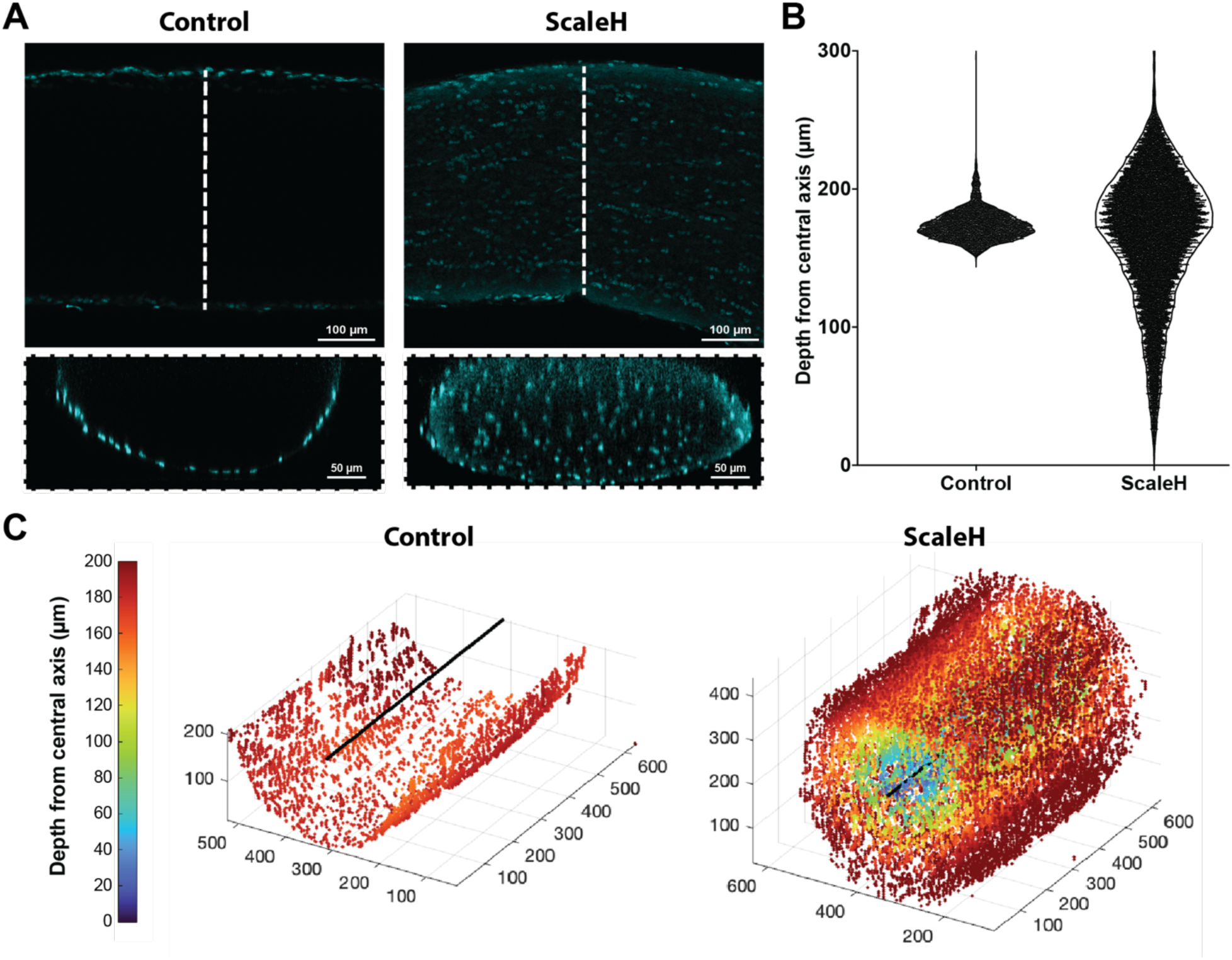
Optical clearing improves nuclear imaging depth and spatial resolution in optic nerves. **(A)** Representative en face and orthographic immunofluorescence images of DAPI- labeled cell nuclei in mouse optic nerves, comparing ProLong Glass (non-cleared control) to ScaleH-cleared samples. Insets show improved nuclear resolution and depth visualization in ScaleH-treated tissue. **(B)** Violin plot showing depth distribution of nuclear fluorescence signals, with ScaleH-cleared nerves exhibiting deeper and broader signal penetration. **(C)** 3D reconstructions of DAPI-labeled nuclei in control and ScaleH-cleared optic nerves, color-coded by depth from the central axis, highlight improved tissue transparency and signal uniformity with ScaleH.

To quantitatively assess signal depth, we analyzed the distribution of nuclear fluorescence along the z-axis. Violin plots of DAPI-labeled cell nuclei demonstrated that ScaleH-cleared nerves exhibited a broader and deeper spread of fluorescence compared to non-cleared controls (Fig. 4B). Three-dimensional reconstructions of DAPI-labeled nuclei, color-coded by depth, further revealed consistent nuclear labeling across the full volume of ScaleH-treated optic nerves, while signal in ProLong Glass samples declined rapidly with depth (Fig. 4C). These results indicate that ScaleH enhances optical clarity and fluorescence signal retention in optic nerve tissue, supporting its effectiveness for whole-mount imaging applications across multiple ocular structures.

## Discussion

Tissue clearing has been widely adopted across various organ systems, prompting laboratories to optimize protocols based on transparency, ease of use, speed, and fluorescence preservation (Kim et al., 2018). In this study, we systematically evaluated five clearing protocols — ScaleS, FOCM, OptiMuS, MACS, and Visikol HISTO-1 — based on their performance in clearing retinal tissues. Our comparison criteria included tissue transparency, color removal, antibody compatibility, and fluorescence retention. Due to the distinct mechanisms by which clearing agents function and the inherent properties unique to retinal tissues, selecting an optimal clearing protocol is critical.

Among the tested protocols, we identified ScaleS as the most effective clearing solution, showing superior performance across nearly all criteria. To further enhance the utility of ScaleS by limiting fluorophore degradation caused by light exposure, oxygen, or prolonged storage (Piña et al., 2022), we developed a novel clearing solution, ScaleH, by incorporating polyvinyl alcohol into the ScaleS formulation. By combining 15% (w/v) PVA and 15% (w/v) glycerol in deionized water, ScaleH self-hardens around the sample, thus mitigating oxygen infiltration and preserving fluorescence signals for extended periods (Gray & Wess, 1950). Importantly, ScaleH retains the user-friendly nature of ScaleS and does not introduce additional steps or complexity for generating cleared and hardened retinal samples (Fig. 3A). The compatibility of ScaleH with both immunostaining and endogenous fluorescence further strengthens its utility for comprehensive structural and cellular analysis of ocular and central nervous system tissues.

In contrast, the two commercial products tested, Visikol HISTO-1 and MACS, had notable limitations. Visikol HISTO-1, an organic solvent-based system, required extensive dehydration steps and introduced tissue fragility and fluorescence degradation despite marginal improvements in transparency over PBS control (Villani et al., 2013). Similarly, MACS, an aqueous-based system employing MXDA instead of urea, also required lengthy dehydration steps that reduced fluorescence signals and increased tissue fragility. Neither commercial option proved advantageous over simpler, in-house solutions for routine retinal applications.

Among the other in-house methods, FOCM and OptiMuS, both aqueous-based, sought to simplify the clearing process while minimizing costs. FOCM, which utilizes DMSO as a polar solvent, achieves substantial tissue transparency but results in significant fluorescence loss, thereby limiting its utility for applications involving antibody labeling or fluorescence microscopy. Conversely, OptiMuS provided better fluorescence preservation, albeit with reduced transparency compared to ScaleS, making it a viable secondary choice.

The success of ScaleS can be attributed to its streamlined, two-step approach, which utilizes ScaleSQ(5) and ScaleS4, along with its excellent fluorescence retention and low cost. These qualities make it particularly well-suited for retinal immunohistochemistry and cell transplantation studies. Our findings underscore its utility in enhancing the visualization and spatial mapping of transplanted RGCs, a critical need in the field given the difficulty of tracking donor cells with limited survival and integration (Oswald et al., 2021)(Ueda et al., 2020)(Zhang et al., 2021)(Soucy et al., 2023). Previous studies have similarly demonstrated the benefits of tissue clearing in visualizing transplanted photoreceptors and axons (Gurdita et al., 2021)(Economo et al., 2019), reinforcing the potential broader applicability of ScaleS and the improved ScaleH in retinal regenerative research.

Additionally, we demonstrated the broader applicability of ScaleH by successfully clearing optic nerve tissue, suggesting its potential for use across other ocular and neural structures. This whole-tissue clearing capability is particularly impactful for studies involving sparse donor axon projections, such as those expected from transplanted RGCs, where traditional sectioning methods risk missing labeled axons altogether. By enabling intact, three-dimensional imaging of the optic nerve, ScaleH allows comprehensive visualization of donor axon trajectories that would otherwise be under-detected or lost in serial sections.

In conclusion, our development of ScaleH significantly advances tissue-clearing methodologies by combining the simplicity and effectiveness of ScaleS with improved long-term fluorescence stability. Its demonstrated efficacy in both retinal and optic nerve tissues suggests strong promise for broader applications in ocular and neurological research. These capabilities will be critical for future studies in regenerative medicine, offering the structural resolution necessary for evaluating therapeutic interventions in the visual system and beyond.

---

**Fig. S1.**
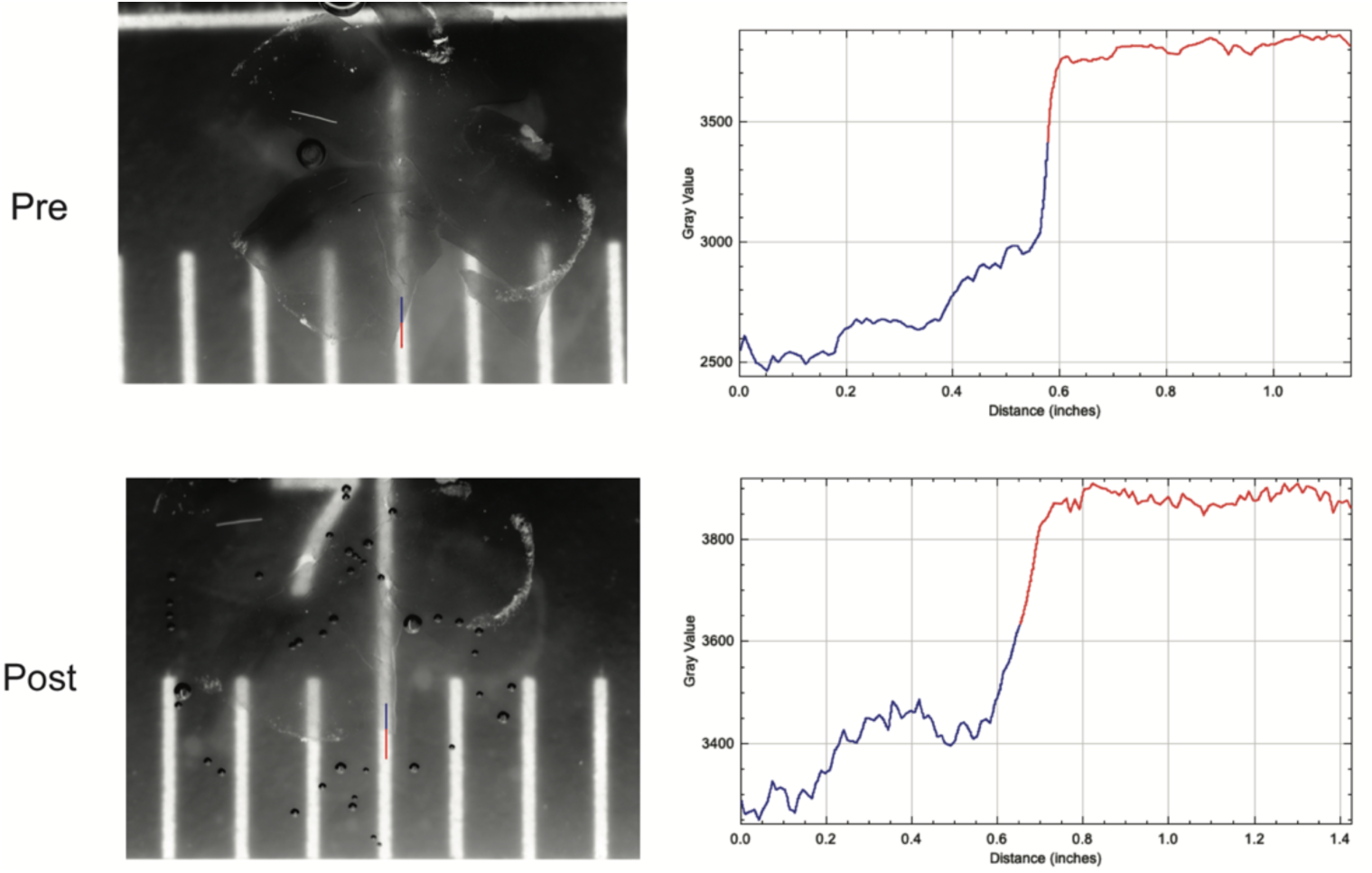
Transparency analysis in pre- and post-cleared retina. **(A)** Representative images of the retina before and after clearing, superimposed on a ruler for analysis. **(B)** Representative ImageJ analysis using the line tool to delineate paths across both unobstructed (red) and retina- obstructed (blue) lines versus intensity gray value.

**Fig. S2.**
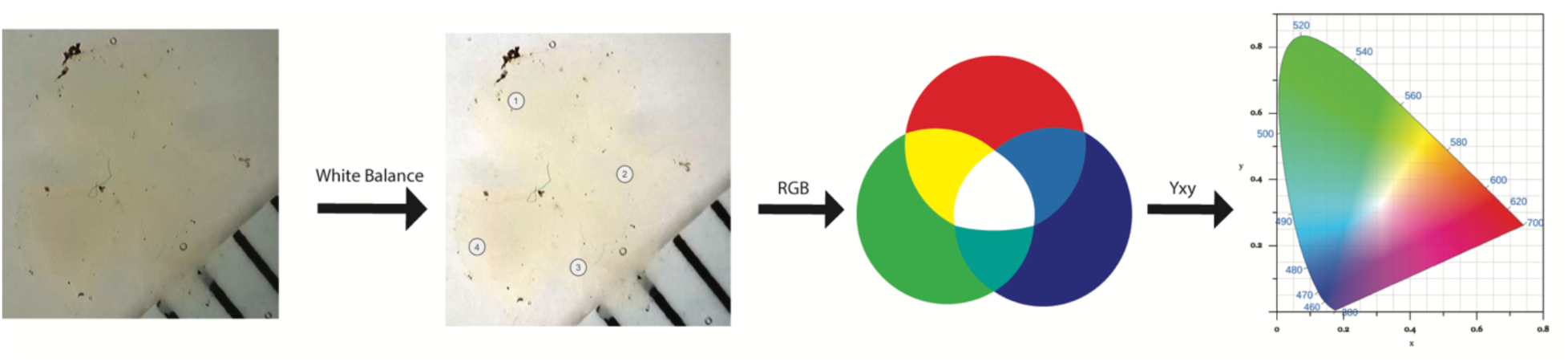
Methodology for color analysis of whole-mount retina. Retina images were white-balanced in Adobe Photoshop, and four points on the extremities of the retina were selected. RGB values were measured in Adobe Photoshop and averaged. RGB values were converted to Yxy values using the EasyRGB value converter and plotted on the CIE 1931 chromaticity diagram.

**Fig. S3.**
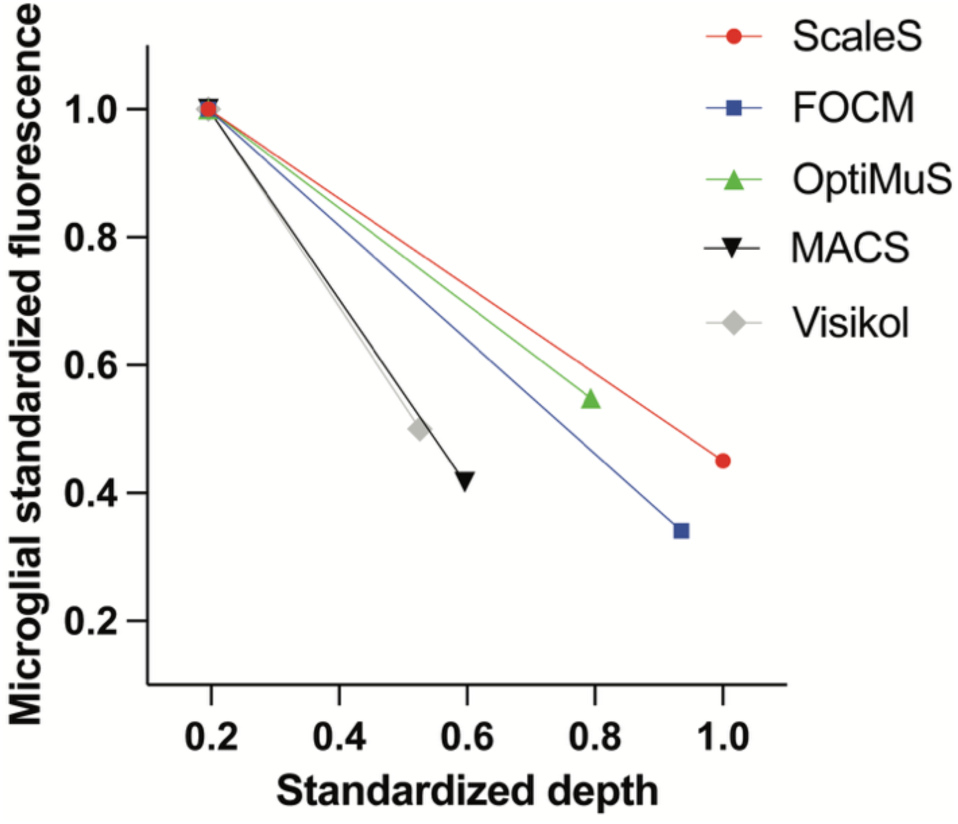
Iba1 slope analysis of microglial layers. Normalized Iba1 intensity plotted against depth for each clearing agent. Slope represents change in fluorescence from superficial to deep microglial layer.

**Fig. S4.**
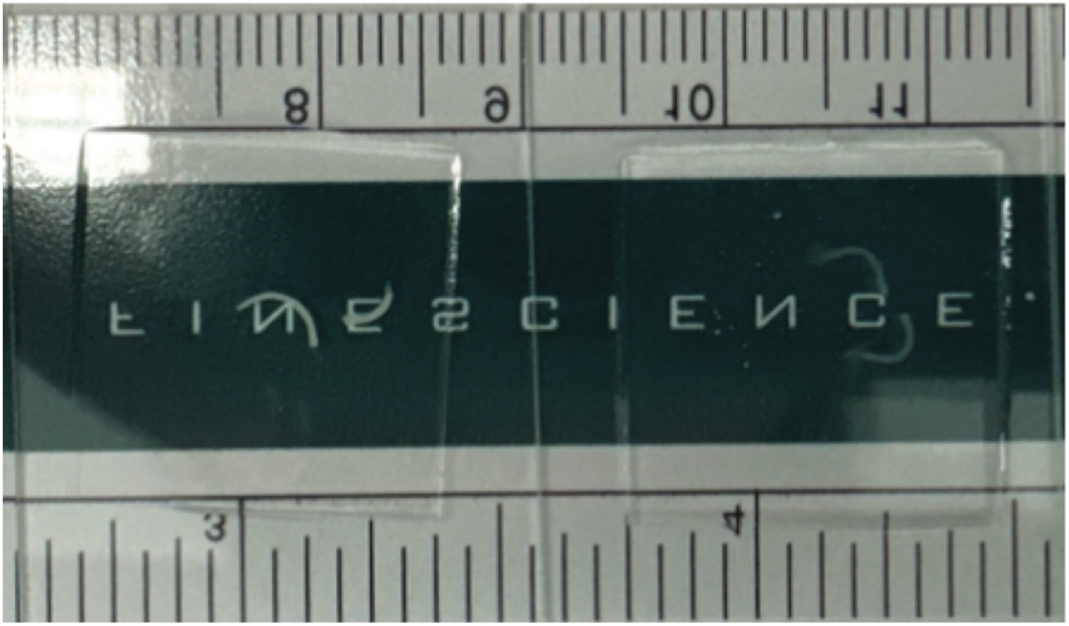
Clearing of mouse optic nerves. Images of optic nerves superimposed on Fine Science Tools ruler. Optic nerves treated with ProLong Glass (non-cleared) on the left and optic nerves treated with ScaleH (cleared) on the right.

**Table S1.** Primary antibodies for immunohistochemistry.

| Antigen | Host | Target Cell / Region | Catalog No. / Supplier | Dilution |
| --- | --- | --- | --- | --- |
| Recoverin | Rabbit | Photoreceptors | ab5585, Chemicon International | 1:500 |
| PKC-α | Mouse | Bipolar cells | sc8393, Santa Cruz Biotechnology | 1:500 |
| Brn3a | Guinea pig | Retinal ganglion cells | 411 004, Synaptic Systems | 1:500 |
| IBA1 | Rabbit | Microglia | ab5076, Abcam | 1:500 |
| PSD-95 | Mouse | Postsynaptic site | N3702-AF647-L, Synaptic Systems | 1:300 |
| VGLUT1 | Rabbit | Glutamatergic synapses | N1602-At488-L, Synaptic Systems | 1:300 |
| mCherry | Chicken | Donor retinal ganglion cells | ab205402, Abcam | 1:400 |

## Notes

### Competing Interest Statement

The authors have declared no competing interest.

### Summary of Updates

Updated the figures and text throughout

### Funding

- National Eye Institute: K99EY036122, F32EY033211, L70EY034355, U24EY029893, P30EY003790
- BrightFocus Foundation: G2020231

## Bibliography

1. Cheng, J., McMahon, S. M., Piston, D. W., & Jackson, M. B. (2023). Comparing confocal and two- photon Ca2+ imaging of thin low-scattering preparations. Biophysical Reports, 3(2), 100109. 10.1016/j.bpr.2023.100109

2. Chung, K., Wallace, J., Kim, S.-Y., Kalyanasundaram, S., Andalman, A. S., Davidson, T. J., Mirzabekov, J. J., Zalocusky, K. A., Mattis, J., Denisin, A. K., Pak, S., Bernstein, H., Ramakrishnan, C., Grosenick, L., Gradinaru, V., & Deisseroth, K. (2013). Structural and molecular interrogation of intact biological systems. Nature, 497(7449), 332–337. 10.1038/nature12107

3. Economo, M. N., Winnubst, J., Bas, E., Ferreira, T. A., & Chandrashekar, J. (2019). Single-neuron axonal reconstruction: The search for a wiring diagram of the brain. Journal of Comparative Neurology, 527(13), 2190–2199. 10.1002/cne.24674

4. Ghoraba, H. H., Akhavanrezayat, A., Karaca, I., Yavari, N., Lajevardi, S., Hwang, J., Regenold, J., Matsumiya, W., Pham, B., Zaidi, M., Mobasserian, A., DongChau, A. T., Or, C., Yasar, C., Mishra, K., Do, D., & Nguyen, Q. D. (2022). Ocular Gene Therapy: A Literature Review with Special Focus on Immune and Inflammatory Responses. Clinical Ophthalmology (Auckland, N.Z.), 16, 1753. 10.2147/OPTH.S364200

5. Graham, H. K., & Duan, X. (2020). Molecular mechanisms regulating synaptic specificity and retinal circuit formation. Wiley Interdisciplinary Reviews. Developmental Biology, 10(1), e379. 10.1002/wdev.379

6. Gray, P., & Wess, G. (1950). The Use of Polyvinyl Alcohol and Its Derivatives as Microscopical Mounting Media. Part I.–Water Miscible Mounting Media. Journal of the Royal Microscopical Society, 70(3), 287–291. 10.1111/j.1365-2818.1950.tb04437.x

7. Gurdita, A., Nickerson, P. E., Pokrajac, N. T., Ortín-Martínez, A., Tsai, E. L. S., Comanita, L., Yan, N. E., Dolati, P., Tachibana, N., Liu, Z. C., Pearson, J. D., Chen, D., Bremner, R., & Wallace, V. A. (2021). InVision: An optimized tissue clearing approach for three-dimensional imaging and analysis of intact rodent eyes. iScience, 24(8), 102905. 10.1016/j.isci.2021.102905

8. Hama, H., Hioki, H., Namiki, K., Hoshida, T., Kurokawa, H., Ishidate, F., Kaneko, T., Akagi, T., Saito, T., Saido, T., & Miyawaki, A. (2015). ScaleS: An optical clearing palette for biological imaging. Nature Neuroscience, 18(10), 1518–1529. 10.1038/nn.4107

9. Hasabeldaim, E. H. H., Swart, H. C., & Kroon, R. E. (2023). Luminescence and stability of Tb doped CaF2 nanoparticles. RSC Advances, 13(8), 5353–5366. 10.1039/D2RA07897J

10. Hashemi, H., Khabazkhoob, Mehdi, Nabovati, Payam, Ostadimoghaddam, Hadi, Shafaee, Shokrolah, Doostdar, Asgar, & and Yekta, A. (2017). The Prevalence of Age-Related Eye Disease in an Elderly Population. Ophthalmic Epidemiology, 24(4), 222–228. 10.1080/09286586.2016.1270335

11. Hoon, M., Okawa, H., Santina, L. D., & Wong, R. O. (2014). Functional Architecture of the Retina: Development and Disease. Progress in Retinal and Eye Research, 42, 44. 10.1016/j.preteyeres.2014.06.003

12. Huang, D., Swanson, E. A., Lin, C. P., Schuman, J. S., Stinson, W. G., Chang, W., Hee, M. R., Flotte, T., Gregory, K., Puliafito, C. A., & Fujimoto, J. G. (1991). Optical Coherence Tomography. *Science (New York*, N.Y.), 254(5035), 1178. 10.1126/science.1957169

13. Jagodzinska, J., Sarzi, E., Cavalier, M., Seveno, M., Baecker, V., Hamel, C., Péquignot, M., & Delettre, C. (2017). Optical Coherence Tomography: Imaging Mouse Retinal Ganglion Cells In Vivo. Journal of Visualized Experiments : JoVE, 127, 55865. 10.3791/55865

14. Kim, J. H., Jang, M. J., Choi, J., Lee, E., Song, K.-D., Cho, J., Kim, K.-T., Cha, H.-J., & Sun, W. (2018). Optimizing tissue-clearing conditions based on analysis of the critical factors affecting tissue-clearing procedures. Scientific Reports, 8(1), 12815. 10.1038/s41598-018-31153-7

15. Kim, K., Na, M., Oh, K., Cho, E., Han, S. S., & Chang, S. (2022). Optimized single-step optical clearing solution for 3D volume imaging of biological structures. Communications Biology, 5(1), 1–10. 10.1038/s42003-022-03388-8

16. London, A., Benhar, I., & Schwartz, M. (2013). The retina as a window to the brain—From eye research to CNS disorders. Nature Reviews Neurology, 9(1), 44–53. 10.1038/nrneurol.2012.227

17. Matsumoto, K., Mitani, T. T., Horiguchi, S. A., Kaneshiro, J., Murakami, T. C., Mano, T., Fujishima, H., Konno, A., Watanabe, T. M., Hirai, H., & Ueda, H. R. (2019). Advanced CUBIC tissue clearing for whole-organ cell profiling. Nature Protocols, 14(12), 3506–3537. 10.1038/s41596-019-0240-9

18. Nanegrungsunk, O., Au, A., Sarraf, D., & Sadda, S. R. (n.d.). New frontiers of retinal therapeutic intervention: A critical analysis of novel approaches. Annals of Medicine, 54(1), 1067–1080. 10.1080/07853890.2022.2066169

19. Oswald, J., Kegeles, E., Minelli, T., Volchkov, P., & Baranov, P. (2021). Transplantation of miPSC/mESC-derived retinal ganglion cells into healthy and glaucomatous retinas. Molecular Therapy. Methods & Clinical Development, 21, 180–198. 10.1016/j.omtm.2021.03.004

20. Piña, R., Santos-Díaz, A. I., Orta-Salazar, E., Aguilar-Vazquez, A. R., Mantellero, C. A., Acosta-Galeana, I., Estrada-Mondragon, A., Prior-Gonzalez, M., Martinez-Cruz, J. I., & Rosas-Arellano, A. (2022). Ten Approaches That Improve Immunostaining: A Review of the Latest Advances for the Optimization of Immunofluorescence. International Journal of Molecular Sciences, 23(3), 1426. 10.3390/ijms23031426

21. Prahst, C., Ashrafzadeh, P., Mead, T., Figueiredo, A., Chang, K., Richardson, D., Venkaraman, L., Richards, M., Russo, A. M., Harrington, K., Ouarné, M., Pena, A., Chen, D. F., Claesson-Welsh, L., Cho, K.-S., Franco, C. A., & Bentley, K. (2020). Mouse retinal cell behaviour in space and time using light sheet fluorescence microscopy. eLife, 9, e49779. 10.7554/eLife.49779

22. Ramirez, A. I., Hoz, R. de, Salobrar-Garcia, E., Salazar, J. J., Rojas, B., Ajoy, D., López-Cuenca, I., Rojas, P., Triviño, A., & Ramírez, J. M. (2017). The Role of Microglia in Retinal Neurodegeneration: Alzheimer’s Disease, Parkinson, and Glaucoma. Frontiers in Aging Neuroscience, 9, 214. 10.3389/fnagi.2017.00214

23. Renier, N., Wu, Z., Simon, D. J., Yang, J., Ariel, P., & Tessier-Lavigne, M. (2014). iDISCO: A Simple, Rapid Method to Immunolabel Large Tissue Samples for Volume Imaging. Cell, 159(4), 896–910. 10.1016/j.cell.2014.10.010

24. Richardson, D. S., & Lichtman, J. W. (2015). Clarifying Tissue Clearing. Cell, 162(2), 246–257. 10.1016/j.cell.2015.06.067

25. Sluch, V. M., Chamling, X., Liu, M. M., Berlinicke, C. A., Cheng, J., Mitchell, K. L., Welsbie, D. S., & Zack, D. J. (2017). Enhanced Stem Cell Differentiation and Immunopurification of Genome Engineered Human Retinal Ganglion Cells. Stem Cells Translational Medicine, 6(11), 1972–1986. 10.1002/sctm.17-0059

26. Soucy, J. R., Aguzzi, E. A., Cho, J., Gilhooley, M. J., Keuthan, C., Luo, Z., Monavarfeshani, A., Saleem, M. A., Wang, X.-W., Wohlschlegel, J., Fouda, A. Y., Ashok, A., Moshiri, A., Chedotal, A., Reed, A. A., Askary, A., Su, A.-J. A., La Torre, A., Jalligampala, A., … The RReSTORe Consortium. (2023). Retinal ganglion cell repopulation for vision restoration in optic neuropathy: A roadmap from the RReSTORe Consortium. Molecular Neurodegeneration, 18(1), 64. 10.1186/s13024-023-00655-y

27. Soucy, J. R., Todd, L., Kriukov, E., Phay, M., Malechka, V. V., Rivera, J. D., Reh, T. A., & Baranov, P. (2023). Controlling donor and newborn neuron migration and maturation in the eye through microenvironment engineering. Proceedings of the National Academy of Sciences, 120(46), e2302089120. 10.1073/pnas.2302089120

28. Tomer, R., Ye, L., Hsueh, B., & Deisseroth, K. (2014). Advanced CLARITY for rapid and high- resolution imaging of intact tissues. Nature Protocols, 9(7), 1682–1697. 10.1038/nprot.2014.123

29. Ueda, H. R., Ertürk, A., Chung, K., Gradinaru, V., Chédotal, A., Tomancak, P., & Keller, P. J. (2020). Tissue clearing and its applications in neuroscience. Nature Reviews. Neuroscience, 21(2), 61–79. 10.1038/s41583-019-0250-1

30. Villani, T. S., Koroch, A. R., & Simon, J. E. (2013). An improved clearing and mounting solution to replace chloral hydrate in microscopic applications. Applications in Plant Sciences, 1(5), 1300016. 10.3732/apps.1300016

31. Zhang, K. Y., Aguzzi, E. A., & Johnson, T. V. (2021). Retinal Ganglion Cell Transplantation: Approaches for Overcoming Challenges to Functional Integration. Cells, 10(6), 1426. 10.3390/cells10061426

32. Zhu, J., Ma, Y., Xu, J., Li, Y., Wan, P., Qi, Y., Yu, T., & Zhu, D. (2021). Dec-DISCO: Decolorization DISCO clearing for seeing through the biological architectures of heme-rich organs. Biomedical Optics Express, 12(9), 5499–5513. 10.1364/BOE.431397

33. Zhu, J., Yu, T., Li, Y., Xu, J., Qi, Y., Yao, Y., Ma, Y., Wan, P., Chen, Z., Li, X., Gong, H., Luo, Q., & Zhu, D. (2020). MACS: Rapid Aqueous Clearing System for 3D Mapping of Intact Organs. Advanced Science, 7(8), 1903185. 10.1002/advs.201903185

34. Zhu, X., Huang, L., Zheng, Y., Song, Y., Xu, Q., Wang, J., Si, K., Duan, S., & Gong, W. (2019). Ultrafast optical clearing method for three-dimensional imaging with cellular resolution. Proceedings of the National Academy of Sciences, 116(23), 11480–11489. 10.1073/pnas.1819583116

